# Divergent paths to seizure-like events

**DOI:** 10.1101/641530

**Authors:** Neela K. Codadu, Robert Graham, Richard J. Burman, R. Thomas Jackson-Taylor, Joseph V. Raimondo, Andrew. J. Trevelyan, R. Ryley Parrish

**Author notes:** Corresponding authors: R Ryley Parrish, Andrew J Trevelyan.

## Abstract

**Aim:** Much debate exists about how the brain transitions into an epileptic seizure. One source of confusion is that there are likely to be critical differences between experimental seizure models. To address this, we compared the evolving activity patterns in two, widely used, *in vitro* models of epileptic discharges.

**Methods:** We compared brain slices, prepared in the same way from young adult mice, that were bathed either in 0 Mg^2+^, or 100µM 4AP, artificial cerebrospinal fluid.

**Results:** We find that while local field potential recordings of epileptiform discharges in the two models appear broadly similar, patch-clamp analysis reveals an important difference in the relative degree of glutamatergic involvement. 4AP affects parvalbumin-expressing interneurons more than other cortical populations, destabilizing their resting state and inducing spontaneous bursting behavior. Consequently, the most prominent pattern of transient discharge (“interictal event”) in this model is almost purely GABAergic, although the transition to seizure-like events (SLEs) involves pyramidal recruitment. In contrast, interictal discharges in 0 Mg^2+^ are only maintained by a very large glutamatergic component that also involves transient discharges of the interneurons. Seizure-like events in 0 Mg^2+^ have significantly higher power in the high gamma frequency band (60-120Hz) than these events do in 4AP, and are greatly delayed in onset by diazepam, unlike 4AP events.

**Conclusions:** The 0 Mg^2+^ and 4AP models display fundamentally different levels of glutamatergic drive, demonstrating how ostensibly similar pathological discharges can arise from different sources. We contend that similar interpretative issues will also be relevant to clinical practice.

## Introduction

Understanding how seizures develop and spread though cortical tissue is of great clinical importance. About 65 million people worldwide are diagnosed as having epilepsy, and many others will experience a seizure at some point in their life ^1,2^. There remain, however, many unresolved issues regarding the basic mechanisms leading to a seizure. Efforts to derive overarching principles regarding epileptic pathophysiology must take into account the key differences between patient phenotypes or between experimental models. Early studies of disinhibited networks showed that, without GABAergic restraint, epileptic discharges could be entrained from single neurons ^3^. This work further provided a detailed, ionic model of the paroxysmal depolarising shift ^4^. The disinhibition models show what can happen with a deficit of GABAergic activity, but cannot address how interneuronal activity might shape epileptic activity. Interneuronal function, on the other hand, has been a dominant theme in studies where epileptic activity is induced either by bathing the tissue in 0 Mg^2+^ ^5–7^ or in 4-aminopyridine (4AP) ^8–10^ ACSF.

In the 0 Mg^2+^ model, the primary cellular effect is to remove the voltage dependent blockade of NMDA receptors^11^, thereby greatly enhancing the postsynaptic effect of any glutamatergic synaptic inputs^11^. In contrast, the primary effect of 4AP is to block certain classes of voltage dependent K^+^ channels. In both cases, the direct pharmacological actions occur rapidly (seconds to minutes), and yet epileptiform activity evolves on a rather slower timescale, over many minutes to hours, suggestive that there are emergent network alterations in each model that are critical for ictogenesis. An added appeal of the 4AP model is that it can be applied *in vivo* ^12–14^, producing ostensibly similar discharges to the *in vitro* activity patterns. Both *in vivo* and *in vitro* 4AP models have been used to good effect in tandem with optogenetic stimulation of specific subpopulations of neurons, indicating how in this special case, interneurons may drive ictogenesis ^15–18^. This is hypothesized to happen through a sequence involving chloride-loading of pyramidal cells and secondary rises in extracellular K^+^ via the potassium chloride cotransporter (KCC2) ^19–22^.

In contrast, the 0 Mg^2+^ model can only be used in vitro (it is harder to remove ions than it is to add a drug), but it, too, involves intense bursts of interneuronal activity ^23–25^, and evidence of short-term changes in GABAergic function ^26,27^. Although, unlike the 4AP model, the interneuronal activity initially appears to act as a restraint on the spreading epileptiform discharges ^28,29^. It is important to resolve these two contrasting views of the role of the interneuronal bursting. We hypothesized that the explanation may lie in the level of concurrent glutamatergic drive in the two models. We therefore made a detailed comparison of activity patterns in these two models, using brain slices prepared in the same way from young adult mice.

## Methods

### Ethical Approval

All animal handling and experimentation were done according to the guidelines laid by the UK Home Office and Animals (Scientific Procedures) Act 1986 and approved by the Newcastle University Animal Welfare and Ethical Review Body (AWERB # 545).

### Slice preparation

Male and female Emx1-Cre (B6.129S2-Emx1tm1(cre)Krj/J; Jackson Laboratory stock number 5628), PV-Cre (B6;129P2-Pvalb<tm1(cre)Arbr>/J; Jackson Laboratory stock number 8069), and SOM-Cre (B6N.Cg.Ssttm2.1(cre)Zjh/J; Jackson Laboratory stock number 18973) mice and C57/B6 mice (ages 3 – 12 weeks) were used in this study. Transgenic mice were back-crossed with the C57/B6 line maintained at Newcastle University, and subsequently maintained on this C57/B6 background (Jackson Laboratory stock number 000664). Mice were housed in individually ventilated cages in a 12 hours light, 12 hours dark lighting regime. Animals received food and water *ad libitum*. Mice were sacrificed by cervical dislocation, brains removed and stored in cold cutting solution (in mM): 3 MgCl_2_; 126 NaCl; 26 NaHCO_3_; 3.5 KCl; 1.26 NaH_2_PO_4_; 10 glucose. For local field potential (LFP) recordings, 400µm horizontal sections were made, using a Leica VT1200 vibratome (Nussloch, Germany). Slices were then transferred to an interface holding chamber and incubated for 1-2 hours at room temperature in artificial CSF (ACSF) containing (in mM): 2 CaCl_2_; 1 MgCl_2_; 126 NaCl; 26 NaHCO_3_; 3.5 KCl; 1.26 NaH_2_PO_4_; 10 glucose. For patch clamp experiments, coronal sections were made at 350µm, and stored in a submerged holding chamber for 1–4 hours prior to experimentation. All the solutions were bubbled continuously to saturate with carboxygen (95% O_2_ and 5% CO_2_).

### Extracellular field recordings

were performed using interface recording chambers. Slices were placed in the recording chamber perfused with modified artificial cerebrospinal fluid (ACSF) to induce epileptiform activity (0 Mg^2+^ or 100µM 4-aminopyrimidine). Recordings were obtained using normal ACSF-filled 1-3MΩ borosilicate glass microelectrodes (GC120TF-10; Harvard Apparatus, Cambridge, UK) placed in deep layers of neocortex. Experiments were performed at 33-36°C. The solutions were perfused at the rate of 3.5mls/min. Waveform signals were acquired using BMA-931 biopotential amplifier (Dataq Instruments, Akron, OH, USA), Micro 1401-3 ADC board (Cambridge Electronic Design, Cambridge, UK) and Spike2 software (v7.10, Cambridge Electronic Design). Signals were sampled at 10kHz, amplified (gain: 500) and bandpass filtered (1-3000Hz). CED4001-16 Mains Pulser (Cambridge Electronic Design) was connected to the events input of CED micro 1401-3 ADC board and is used to remove 50Hz hum offline. SLEs were visually identified with their start time as the time of occurrence of high-frequency rhythmic bursts (tonic-phase) associated with high-frequency signals, and the events were considered to end when the interval between two after-discharges (clonic-phase) is ≥ 2s. Frequency analysis of the recordings were performed using a custom-written code in Matlab2018b (MathWorks, Natick, MA, USA).

### Viral injections

PV-Cre and SOM-Cre, or Emx1-Cre pups were injected with AAV5.Syn.Flex.tdTomato, purchased from the University of Pennsylvania vector core. Injections were performed either on the day of birth, or the following day. Pups were set on a stereotaxic frame and anesthetized with isoflurane, following application of EMLA cream (2.5% lidocaine and 2.5% prilocaine) to the left top of their head. Injections were made, using a 10μl Hamilton syringes with a bevelled 36-gauge needle (World Precision Instruments, Sarasota, FL, USA), unilaterally into the lateral ventricle and overlying cortical plate, at about 1 mm anterior to lambda and 1 mm lateral to the midline into the left hemisphere, starting at 1.7 mm deep from the top of the dura mater, for a total of four separate 50nl injections, deepest first and coming up 0.3 mm for each subsequent injection. Approximately 200nl (∼1000 viral particles) were injected into the left hemisphere over a 10 minute period. Pups were monitored until they awoke following the procedure and then returned to their home cage. These neonatal injections produced widespread cortical expression of tdTomato into the neurons of interest.

### Patch-clamp recordings

Slices were perfused at 3-5mls/min and heated to 33-34°C. Whole cell data were acquired using pClamp software, Multiclamp 700B, and Digidata acquisition board (Molecular Devices, Sunnyvale, CA, USA). Whole cell recordings of layer 5 neurons were made using 4-7MΩ pipettes. Pipettes were filled with a KMeSO_4_-based internal solution containing (mM): 125 KMeSO_4_, 6 NaCl, 10 HEPES, 2.5 Mg-ATP, 0.3 Na_2_-GTP. For the voltage clamp recordings, 5mM QX-314 (N-(2,6-dimethylphenylcarbamoylmethyl) was added to the internal solution to block action potential firing while the cells were held at −30mV. Osmolarity and pH of all the internal solutions used were adjusted to 284mOsm and 7.4. The targeted patch experiments of the PV-interneurons, SST-interneurons, and pyramidal neurons were done in current clamp mode, with the cells being identified by the presence of tdTomoto. The fluorescence was visualized using an Olympus DSU spinning disk BX/50WI upright microscope (UMPlanFL N 20x, 0.5 NA objective; Olympus, UK), illuminated using a Mercury arc lamp, controlled with a fast Sutter shutter (Sutter Instrument, USA), using the standard Olympus rhodamine (U-MRFPHQ) filter set. The system utilizes a Hamamatsu C9100 EM camera (Hamamatsu Photonics, Japan) to collect images, run by Simple PCI software (Digital Pixel, UK) installed on Dell Precision computers (Dell, UK). Patch clamp data was analysed using custom-written codes in Matlab2018b. Briefly, epileptiform discharges were identified from the voltage clamp recordings and classified automatically. In short, the events immediately prior to seizures, occurring at ~1-2Hz, and progressing straight into a full SLE, are referred to as “preictal events”. Other transient discharges (<500ms duration) which occur at a much lower rate (~0.1Hz) are referred to as “interictal events” (note that these also include events prior to the first SLE, even though as such, these events are not “inter”-ictal). Trace deflections fitting a generalised model were grouped by their structure as excitatory only, inhibitory only. All events were considered for the composite category and were recategorised only if they displayed bidirectional deflections with respect to a dynamically calculated baseline.

### Pharmacology

Drugs acting at glutamatergic and GABAergic receptors were used at the following concentrations: 50µM D-2-amino-5-phosphonovalerate (AP5); 20µM NBQX disodium salt (Hello Bio, Bristol, UK); 10μM Gabazine (Hello Bio, Bristol, UK); 3µM Diazepam (Tocris, Abingdon, UK); 5μM CPG-55845.

### Data Analysis and Statistics

Data were analysed offline using Matlab R2018b (MathWorks, USA). Groups of two were analysed using the Mann-Whitney test or Student’s t-test. A Kruskal-Wallis test followed by a Dunn’s multiple comparison test was used for data with 3 or more groups. Proportion data was analysed with a Chi-square test. Significance was set at P ≤ 0.05 for all analyses.

## Results

### Superficial similarities between 0 Mg^2+^ and 4AP induced epileptiform activity

Brain slice preparations have long been used to understand how epileptic discharges are manifest in cortical networks. We compared the neocortical activity patterns that occur in two widely used models, induced by bathing brain slices in either 0 Mg^2+^ or 100µM 4-aminopyridine (4AP) artificial cerebrospinal fluid (ACSF). Both models induce an evolving pattern of interictal and seizure-like activity, in neocortex, eventually progressing into a pattern of rhythmic epileptiform discharges, termed late recurrent discharges (LRDs), that has been likened to status epilepticus (Figure 1Ai, Aii) ^6,30^. The time to the first full ictal event and time to the LRD in 4AP is significantly shorter than in 0 Mg^2+^ but otherwise, the activity patterns appear broadly similar (Figure 1B-E).

**Figure 1.**
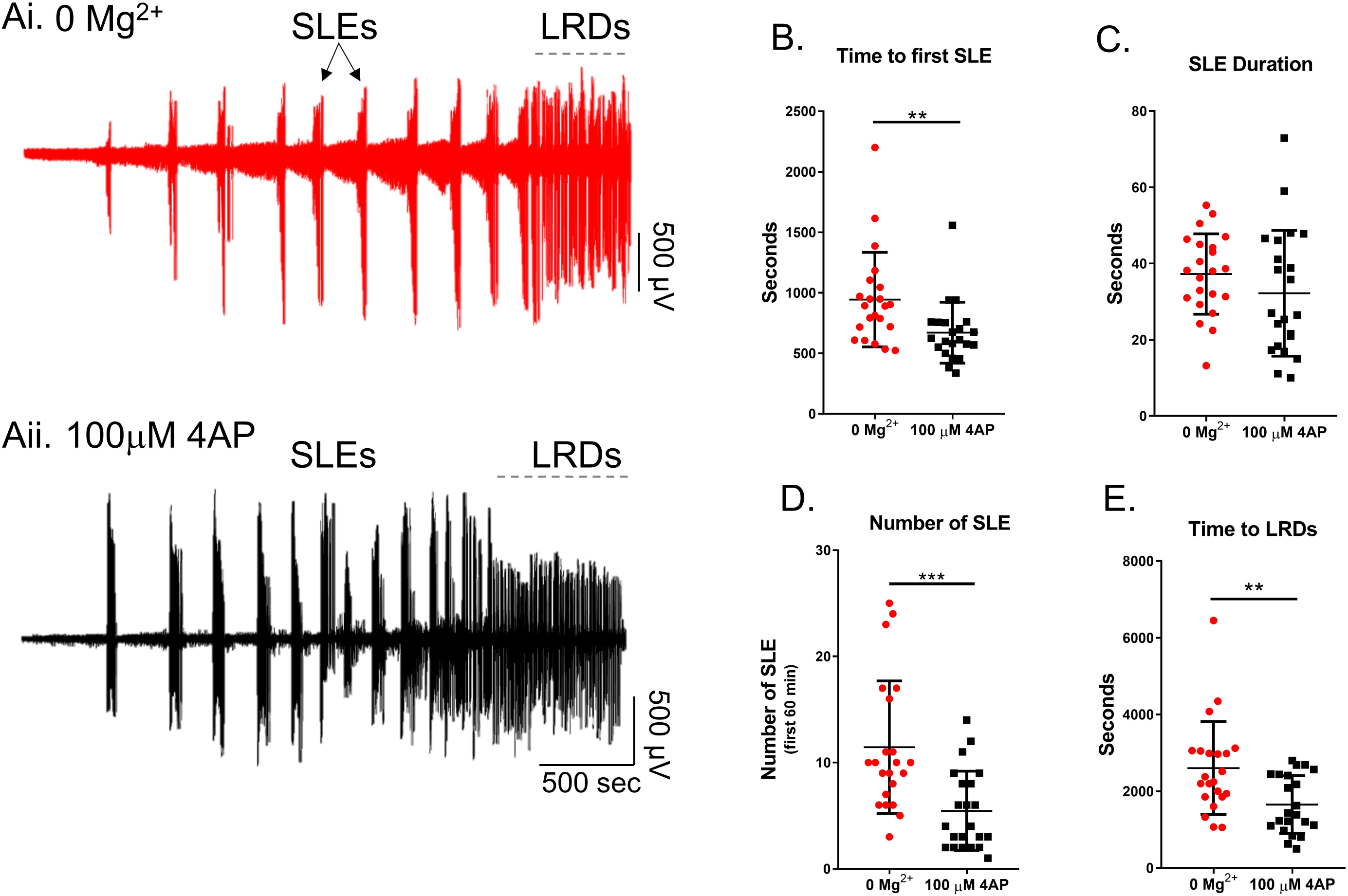
Similarities between evolving epileptiform activity, in 0 Mg^2+^ and 4AP ACSF. Ai) Example trace from 0 Mg^2+^ induced epileptiform activity. Aii) Equivalent recording in 4AP. B) The 4AP model leads to SLE significantly faster than the 0 Mg^2+^ model (Mann-Whitney test, **p = 0.0017). C) There is no difference between the duration of seizures between the 4AP and 0 Mg^2+^ models (Mann-Whitney test, p = 0.1733). D) There is no difference in the number of SLE between the 4AP and 0 Mg^2+^ models (Mann-Whitney test, ***p = 0.0002). E) The 4AP model enters LRD significantly earlier than the 0 Mg^2+^ model (Mann-Whitney test, **p = 0.0050).

Extracellular recordings, however, are rather abstracted representations of the underlying activity, and so while they clearly have a large informational content, the underlying neuronal causes of the signal often remains rather cryptic. We therefore explored the two models using patch-clamp recordings of pyramidal cells (Figure 2). Large, layer 5 pyramidal cells have dendritic trees that extend through the full depth of the cortex, meaning that recordings from these cells provide a good sampling of synaptic drives onto a cortical column. We recorded 19 pyramidal cells (10 cells in 0 Mg^2+^; 9 cells in 4AP) in voltage clamp, holding the cells at −30mV, roughly halfway between the reversal potential of glutamatergic and GABAergic receptors. This approach had previously proved highly informative about the pattern of synaptic bombardment during spreading ictal events^23^. In 4AP, most early interictal-like discharges were manifest as entirely upward deflections, suggestive of an overwhelming preponderance of GABAergic activity. In contrast, the majority of 0 Mg^2+^ events showed prominent downward deflections, indicative of large glutamatergic drive. The two models were highly significantly different (Figure 2Aii, Bii). These same patterns were replicated in the events immediately prior to ictal events (pre-ictal activity; Figure 3Ai, Bi), and which likewise showed a highly significant difference in the level of glutamatergic drive in the two models (Figure 3Aii, Bii). Taken together, these data demonstrate that despite the ostensible LFP similarities, there are important differences between the models, particularly in the pattern of synaptic drive onto the pyramidal cell population, suggesting that different cell populations may drive the generation of pathological discharges in the two models.

**Figure 2.**
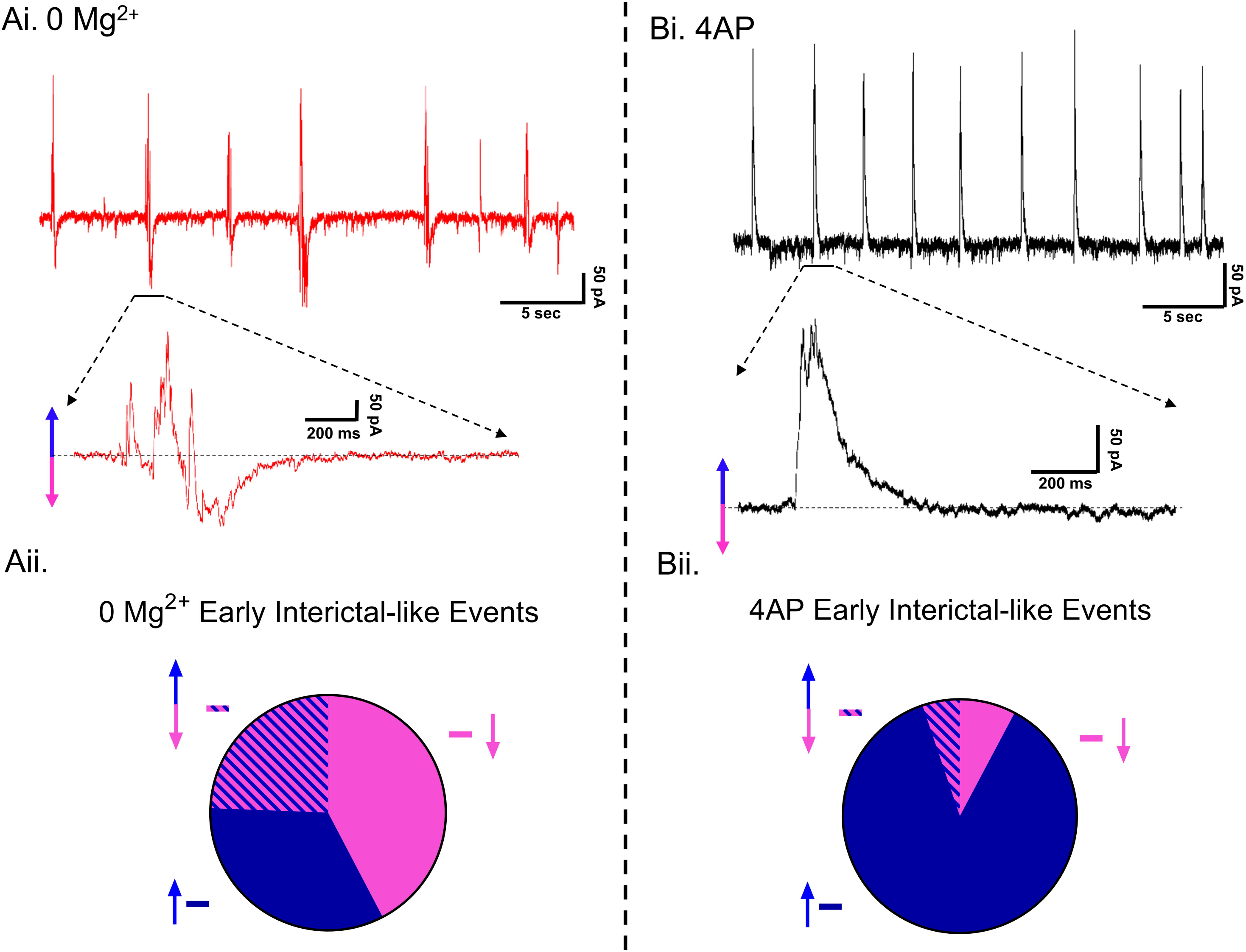
Different levels of glutamatergic drive in early interictal-like events, in the two models. Ai) Example patch-clamp recording of typical early interictal-like events onto a layer 5 pyramidal cell induced by washout of Mg^2+^ ions (Cell is being held at −30 mV). Aii) Pie chart demonstrating the proportion of putative IPSCs (33.3%, represented with an upward blue arrow), EPSCs (42.3%, represented with a downward pink arrow), and composite events (events containing both a putative IPSC and an EPSC; 24.4%, represented with both an upward blue arrow and a downward pink arrow) of early interictal-like postsynaptic currents onto pyramidal cells after washout of Mg^2+^ ions (4,002 events were analysed from 10 slices). Bi) Equivalent recording in 4AP as shown in Ai (Cell is being held at −30 mV). Bii) Pie chart demonstrating the proportion of putative IPSCs (87.0%, represented with an upward blue arrow), EPSCs (7.8%, represented with a downward pink arrow), and composite events (5.2%, represented with both an upward blue arrow and a downward pink arrow) of early interictal-like postsynaptic currents onto pyramidal cells after addition of 4AP (9,992 events were analysed from 9 slices, Proportion data from 4AP is significantly different from the 0 Mg^2+^ paradigm, χ^2^ test, p < 0.0001).

**Figure 3.**
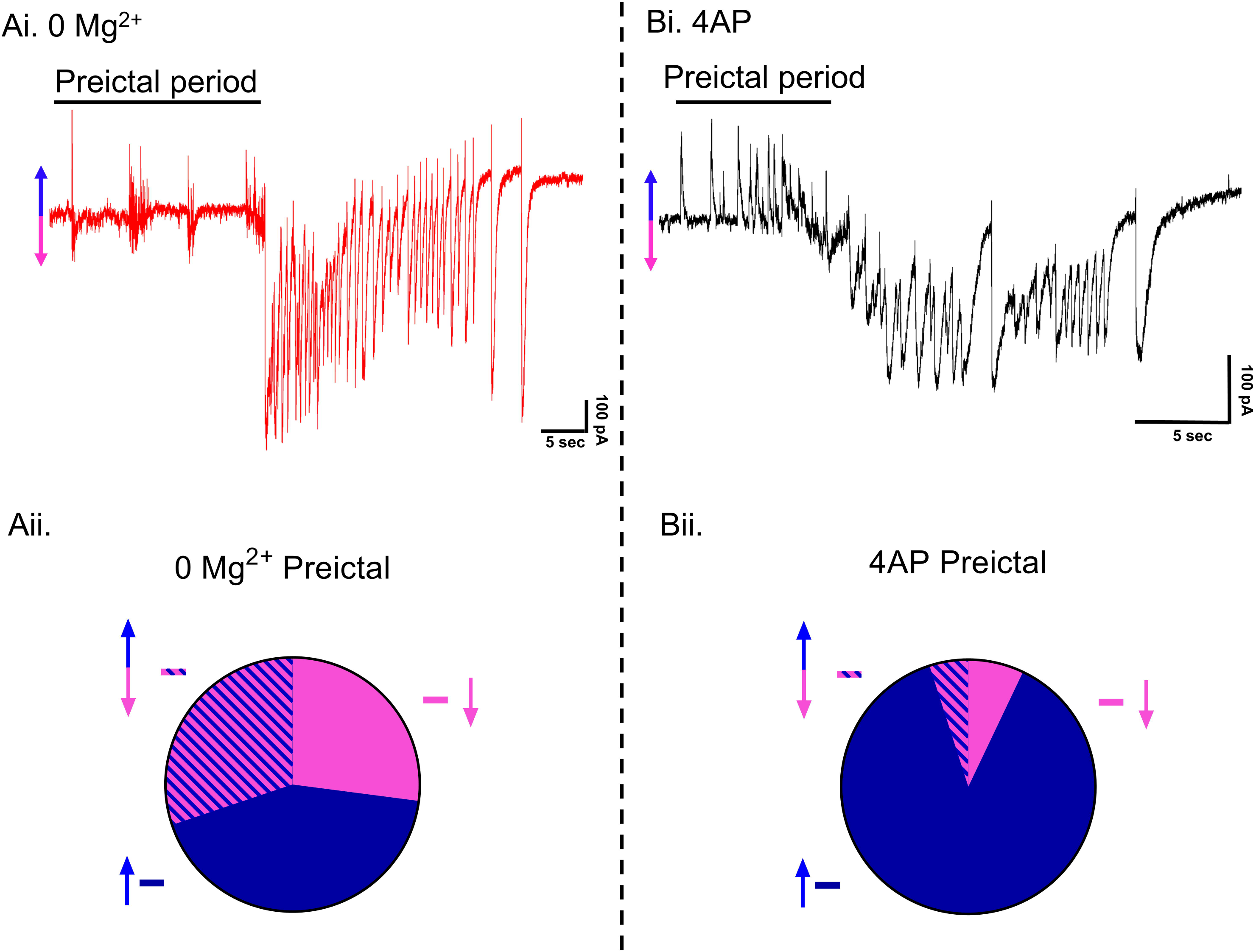
Different levels of glutamatergic drive during preictal activity, in the two models. Ai) Example patch-clamp recording of typical SLE induced by washout of Mg^2+^ ions (Cell is being held at −30 mV). Aii) Pie chart demonstrating the proportion of putative IPSCs (42.8%, represented with an upward blue arrow), EPSCs (27.0%, represented with a downward pink arrow), and composite events (events containing both a putative IPSC and an EPSC; 30.2%, represented with both an upward blue arrow and a downward pink arrow) of preictal postsynaptic currents onto pyramidal cells after washout of Mg^2+^ (152 events were analysed from 10 slices). Bi) Equivalent recordings in 4AP as shown in Ai (Cell is being held at −30 mV). Bii) Pie chart demonstrating the proportion of putative IPSCs (88.0%, represented with an upward blue arrow), EPSCs (7.0%, represented with a downward pink arrow), and composite events (5.0%, represented with both an upward blue arrow and a downward pink arrow) of preictal postsynaptic currents onto pyramidal cells after addition of 4AP (105 events were analysed from 9 slices, Proportion data from 4AP is significantly different from the 0 Mg^2+^ paradigm, χ^2^ test, p < 0.0001).

We next examined the source of the ictal-like activity through pharmacological dissection of the glutamatergic and GABAergic components of the network. As expected, application of AMPA and NMDA receptor blockers resulted in the abolishment of all interictal and ictal-like activity in the 0 Mg^2+^ model (Figure 4A). In contrast, in the 4AP model, application of glutamate receptor blockers only blocked the ictal-like activity, but interictal activity persisted, as has been previously reported ^10,31^. This interictal activity was eliminated through the application of the of GABA_A_ receptor blocker gabazine (Figure 4B), strongly suggesting that this remaining epileptiform activity is due to post-synaptic IPSCs onto the pyramidal cells ^10^. This demonstrates that while seizure-like events in both models are downstream of glutamate receptors activation, the source of the interictal activity is fundamentally different between the two models.

**Figure 4.**
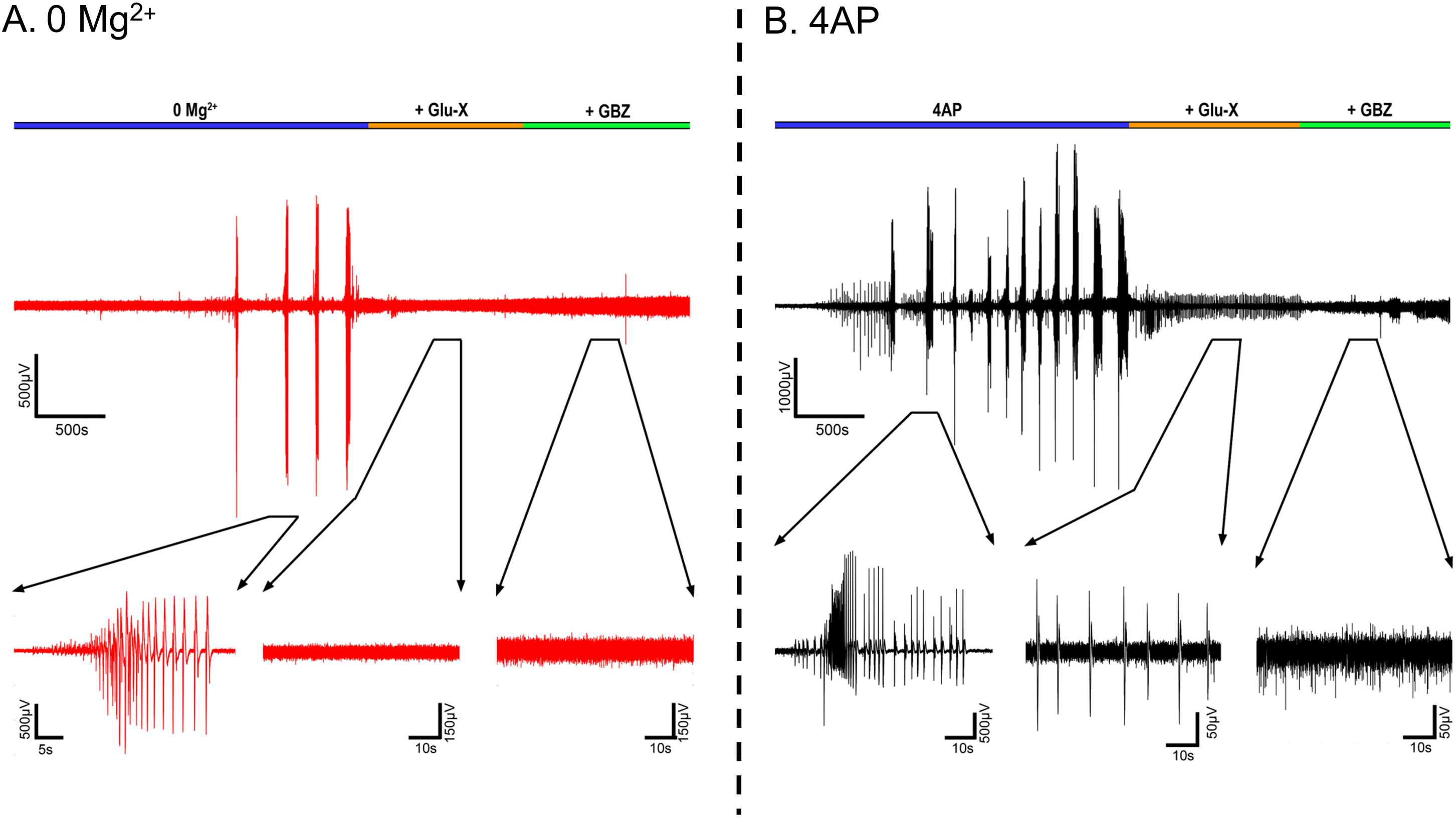
4AP interictal-like activity is sustained in the presence of glutamatergic blockers but not in the 0 Mg^2+^ paradigm. A) Representative LFP trace of a 0 Mg^2+^ recording before and after addition of Glu-X (NBQX and AP5) and Glu-X plus Gabazine (N = 3). B) Equivalent recordings in 4AP (N = 3).

### 4AP preferentially activates PV interneurons

In order to understand the source of the persistent epileptiform activity in the 4AP model, we examined the direct effects of 4AP on firing patterns in the three main classes of neocortical neurons: pyramidal cells, and both parvalbumin-expressing (PV), and somatostatin-expressing (SST) interneurons. Recordings were performed in the absence of any fast synaptic neurotransmission (glutamatergic blockade: 20µM NBQX and 50µM AP5; GABAergic blockade: 10μM gabazine and 5μM CPG-55845). 4AP induced spontaneous action potential firing in all cell classes studied (Figure 5A), but the effect on PV-interneurons was larger than on the other two cell classes (Figure 5B). Consistent with this finding, 4AP application had no appreciable effect on the resting membrane potential, or input resistance, of either pyramidal cells or SST interneurons, but it induced a highly significant increase in the input resistance of PV interneurons (Table 1). Additionally, 4AP reduced action potential threshold in both PV and pyramidal cells (but not SST interneurons), but this only translated into an increase in the evoked firing (triggered by a 100pA current injection) in PV interneurons, but not in the pyramidal cells (Table 1; Figure 5D; 6B). 4AP further caused a highly significant broadening of action potentials in all cell classes, although the effect was much larger in the interneuronal classes (84% and 104% increase in PV and SST interneurons respectively, but only 18% increase for pyramidal cells, Figure 6A;Table 1).

**Figure 5.**
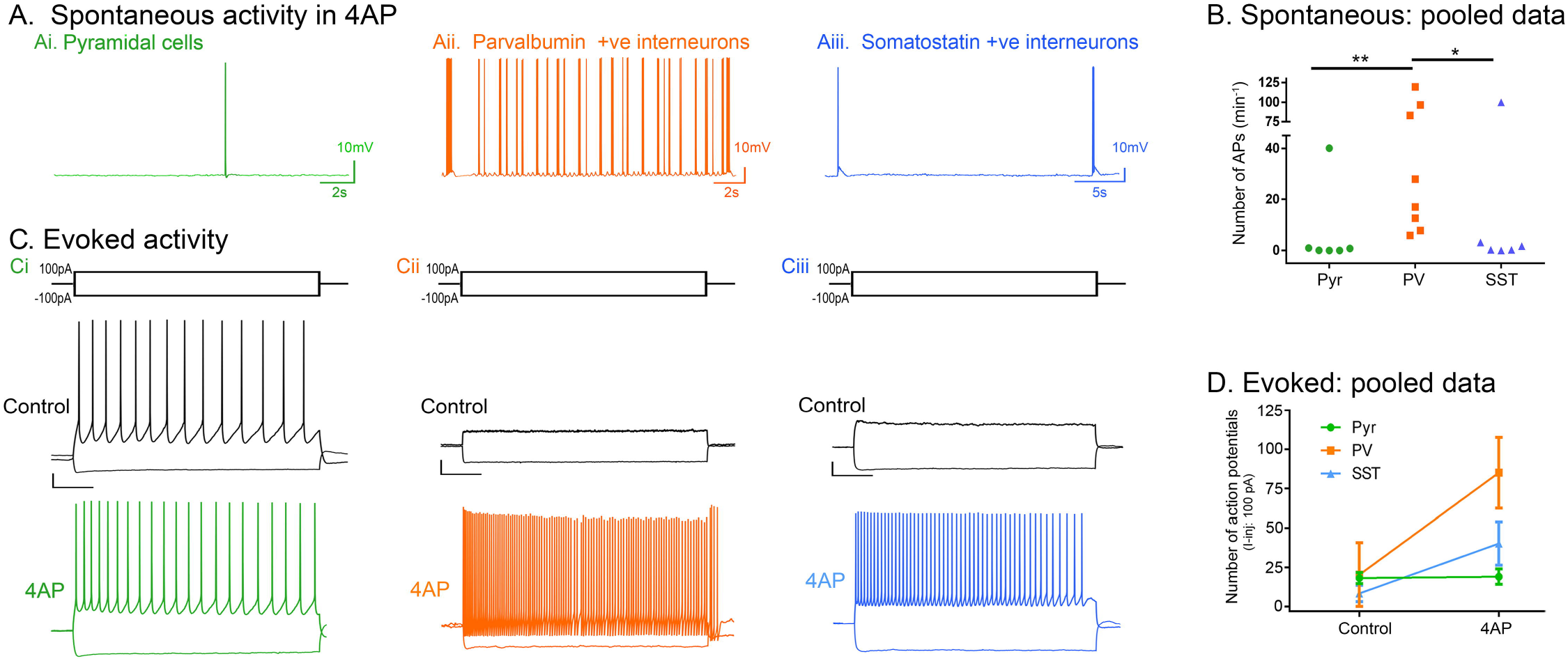
4AP preferentially induces increased action potential firing in the PV cell population compared to pyramidal and SST cells. A) Spontaneous activity: representative traces of a pyramidal cell, a PV-interneuron, and a SST-interneuron following wash in of 100µM 4AP with Glutamate (NBQX and AP5) and GABA (Gabazine and CPG-55845) receptor blockers (N = 6). B) Spontaneous action potential firing in the PV cells is significantly higher than in the pyramidal or SST cells (Kruskal-Wallis Test with post-hoc test, **p = 0.008 for pyramidal vs PV, *p = 0.042 for SST vs PV). C) Evoked activity: representative traces showing responses of Ci) a pyramidal cell, Cii) a PV interneuron, and Ciii) a SST interneuron to 3s long hyperpolarisation (−100pA) and depolarisation (100pA) current pulses, in the presence of 4AP (GABA and glutamatergic neurotransmission blocked). Scale bars: horizontal, 500ms; vertical, 10mV. D) 4AP increased the number of action potentials fired by PV interneurons (Wilcoxon matched-pairs rank test, p = 0.008; N = 9) due to 100pA current injection, but not by pyramidal cells (paired t-test, p = 0.769; N = 6) or SST interneurons (Wilcoxon matched-pairs rank test, p = 0.125; N = 6).

**Figure 6.**
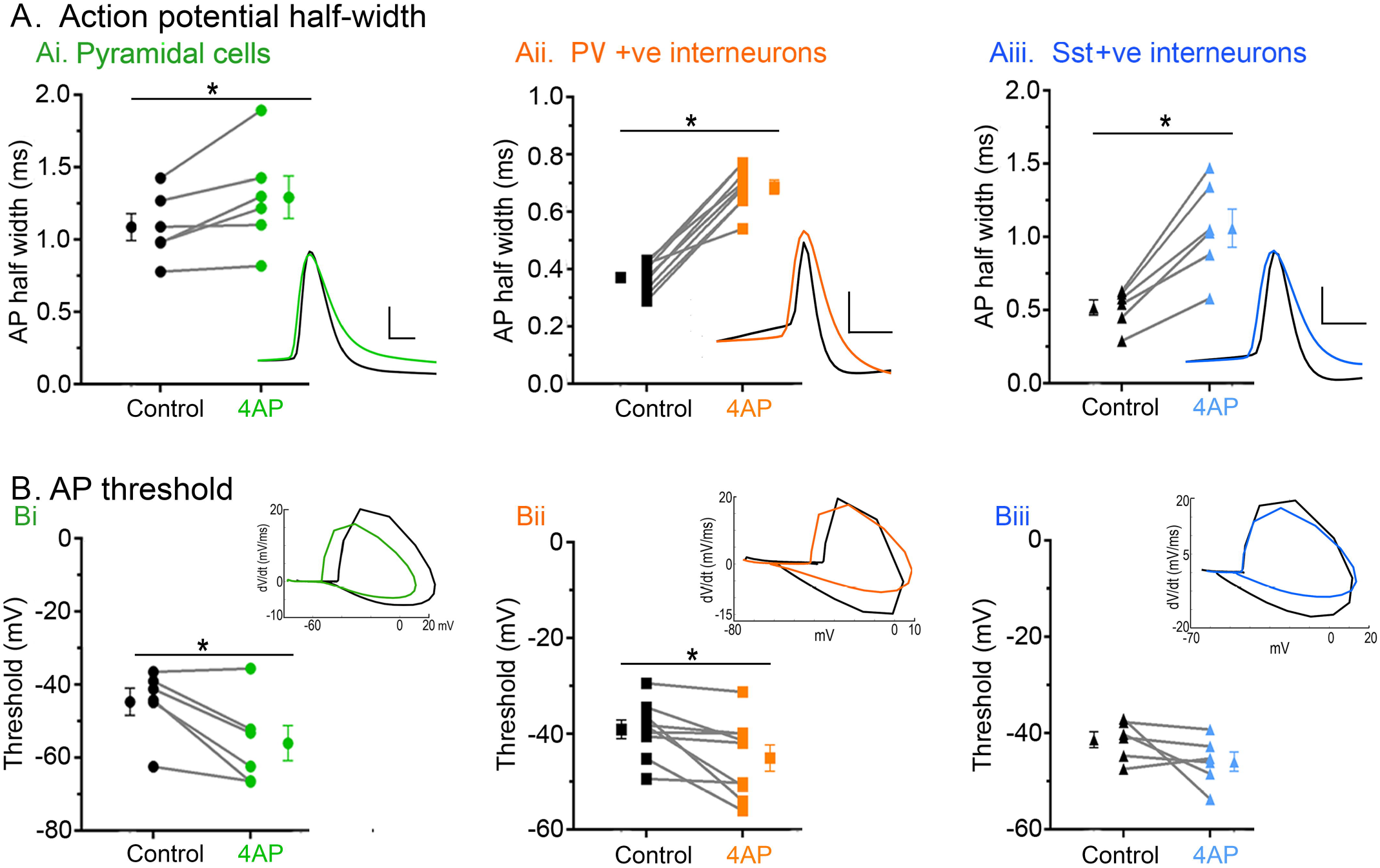
4AP affects the firing properties in both PV- and SST-expressing interneurons, and also pyramidal cells. Ai) 4AP significantly increased the action potential half-width of pyramidal cells (paired t-test, p = 0.033; N = 6). Inset: representative pyramidal cell action potential trace before (black) and after (green) 4AP. Scale bars: horizontal, 1ms; vertical, 20mV. Aii) 4AP significantly increased the action potential half-width of PV interneurons (paired t-test, p < 0.0001; N = 9). Inset: representative PV interneuron action potential trace before (black) and after (orange) 4AP. Scale bars: horizontal, 1ms; vertical, 20mV. Aiii) 4AP significantly increased the action potential half-width of SST interneurons (paired t-test, p = 0.002; N = 6). Inset: representative SST interneuron action potential trace before (black) and after (blue) 4AP. Scale bars: horizontal, 1ms; vertical, 20mV. Bi) 4AP significantly reduced the action potential threshold of pyramidal cells (paired t-test, p = 0.023; N = 6). Inset: representative phase plots of pyramidal cell action potential trace before (black) and after (green) 4AP. Bii) 4AP significantly reduced the action potential threshold of PV interneuron (paired t-test, p = 0.021; N = 9). Inset: representative phase plots of PV interneuron action potential trace before (black) and after (orange) 4AP. Biii) 4AP had no effect on the action potential threshold of SST interneuron (paired t-test, p = 0.159; N = 6). Inset: representative phase plots of SST interneuron action potential trace before (black) and after (blue) 4AP.

**Table 1.**
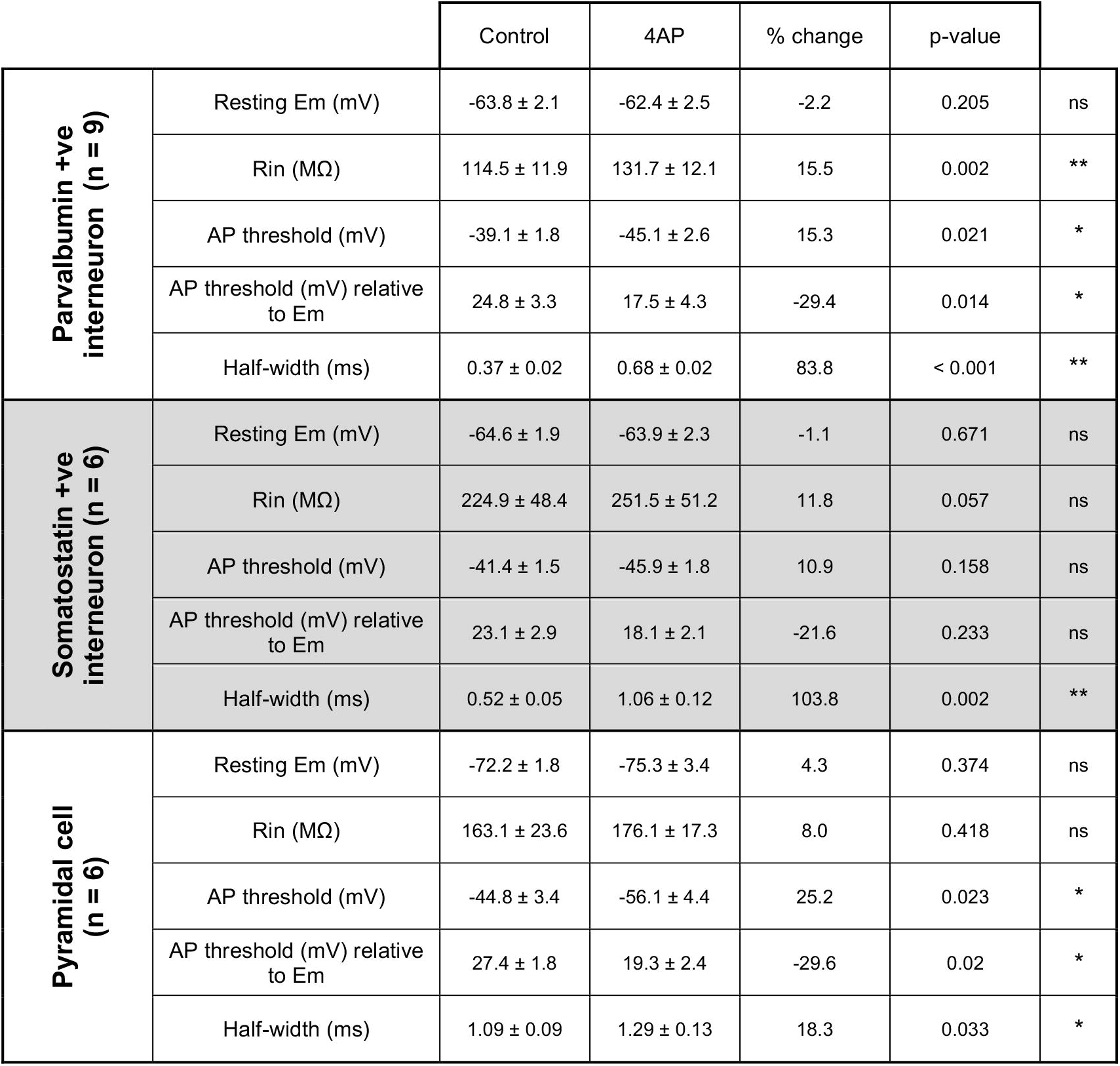

We concluded that 4AP affects PV-interneuron firing more than it does SST and pyramidal firing, leading to a preferential activation of the PV cell class. PV interneurons are, thus, likely to be the main source of the persistent pathological discharges following application of glutamate receptor blockers, in this model. We therefore performed targeted, cell-attached patch clamp recordings of the PV interneuron population during spontaneous interictal events in the two models (no glutamatergic blockade). We found that while the mean and maximal firing rates during these interictal bursts were comparable in the two models, these bursts of firing lasted approximately twice as long in the 4AP events and comprised roughly three times as many action potentials per event (Figure 7).

**Figure 7.**
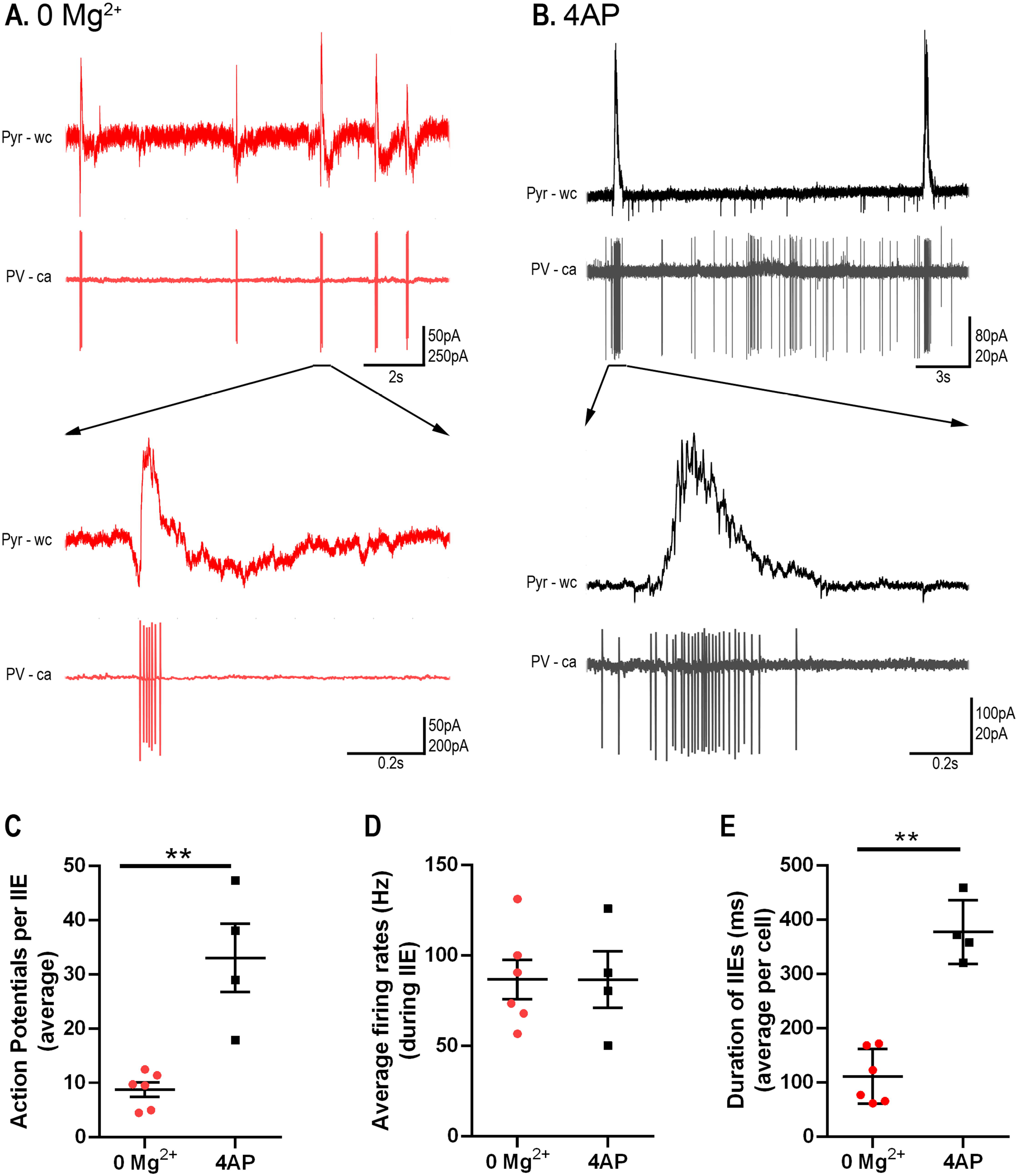
PV firing during interictal events is more sustained in 4AP than in 0 Mg^2+^ events. A) Example paired recordings of pyramidal cells (“Pyr”, recorded in whole cell (“wc”) voltage-clamp mode at −30mV) and PV interneurons (recorded in cell-attached (“ca”) mode) in 0 Mg^2+^. Note the intense bursts of PV firing coincident with upwards deflections of the pyramidal trace, consistent with the PV activity being the source of the GABAergic synaptic barrages. B) Equivalent example paired recordings in 100µM 4AP. C) PV interneurons fire significantly more action potentials per interictal event in 4AP than they do in 0 Mg^2+^ (79 events from 6 slices analysed from the 0 Mg^2+^ paradigm while 57 events from 4 slices were analysed for the 4AP model; Mann-Whitney test, **p = 0.0095). D) There is no difference in the PV cell firing rates during interictal events between the models. E) Duration of interictal events are longer in the 4AP model than in 0 Mg^2+^ model (Mann-Whitney test, **p = 0.0095).

We next analysed prolonged extracellular recordings of 22 brain slices bathed in 4AP and 24 slices bathed in 0 Mg^2+^, to investigate whether the difference in glutamatergic involvement between the models was evident in the local field potential. To control for differences in the electrode quality, we normalised the traces to the maximal field deflection during full ictal events, since we reasoned that these are likely to reflect comparable levels of network activation in the two models. In each model, in almost identical proportions, we found instances of both low voltage, fast onset, (0 Mg^2+^ model 43% of events; 4AP, 44% of events) and hypersynchronous onset (0 Mg^2+^, 57% of events; 4AP, 56% of events). A power spectra analysis over the entire ictal events indicated that there was a significant excess power at frequencies in the high gamma range in the 0 Mg^2+^ (Figure 8). This excess power observed in the 0 Mg^2+^ model extended also into the fast ripple range (250 to 500 Hz, data not shown). Another notable difference between the models is that the post-ictal refractory period lasted far longer in 0 Mg^2+^ (time delay after the first seizure, before the resumption of interictal discharges =106 ± 67s; n = 24 brain slices) compared with that in 4-AP (post-ictal delay = 25 ± 50s; n = 22; p = 3.0e-5).

**Figure 8.**
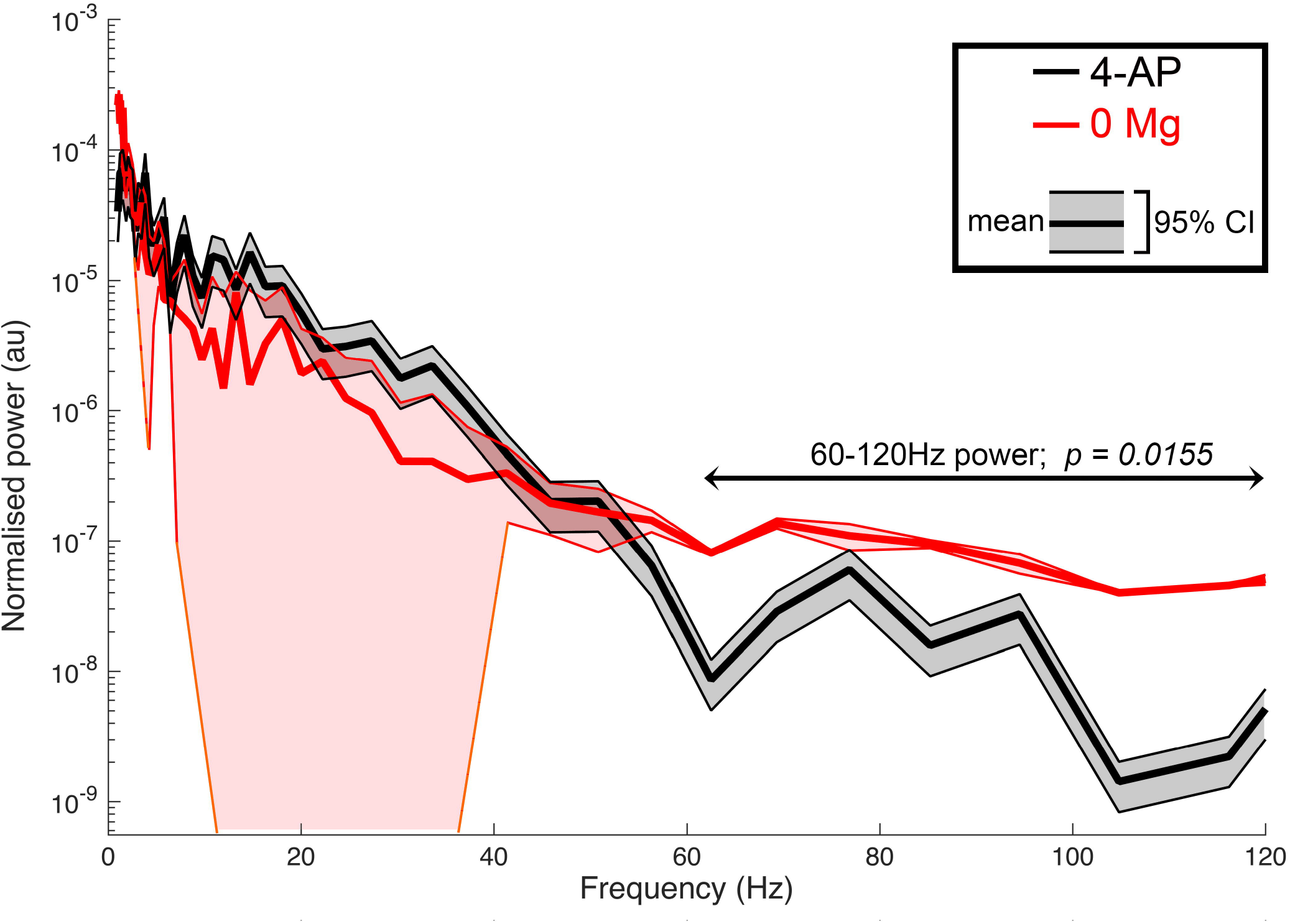
0 Mg^2+^ and 4AP SLEs differ in higher frequency power. 0 Mg^2+^ seizures display more power in the higher frequency range then 4AP seizures (mean summed power 60-120Hz (normalised traces), 0 Mg^2+^ = 7.08e-7 ± 8.99e-7; 4-AP = 1.78e-7 ± 4.21e-7; p = 0.0155; 22 and 24 brain slices respectively).

### The 0 Mg^2+^ and 4AP models differ in their sensitivity to diazepam

Finally, we examined whether there were differences in sensitivity to the benzodiazepine family of drugs between the models as a result of their different drives. We found that bath application of diazepam significantly delayed the onset of the SLE and increased the duration of the early interictal period in the 0 Mg^2+^ model, while having no effect in the 4AP model (Figure 9A-D). Additionally, in the 0 Mg^2+^ model, the number and rate of early interictal-like events were greatly increased by diazepam. In contrast, diazepam affected neither in 4AP (Figure 9E, F). These data further suggest that these models produce epileptiform activity by different mechanisms and serve to illustrate how important it is to understand the source of epileptiform activity, when considering how to manage it.

**Figure 9.**
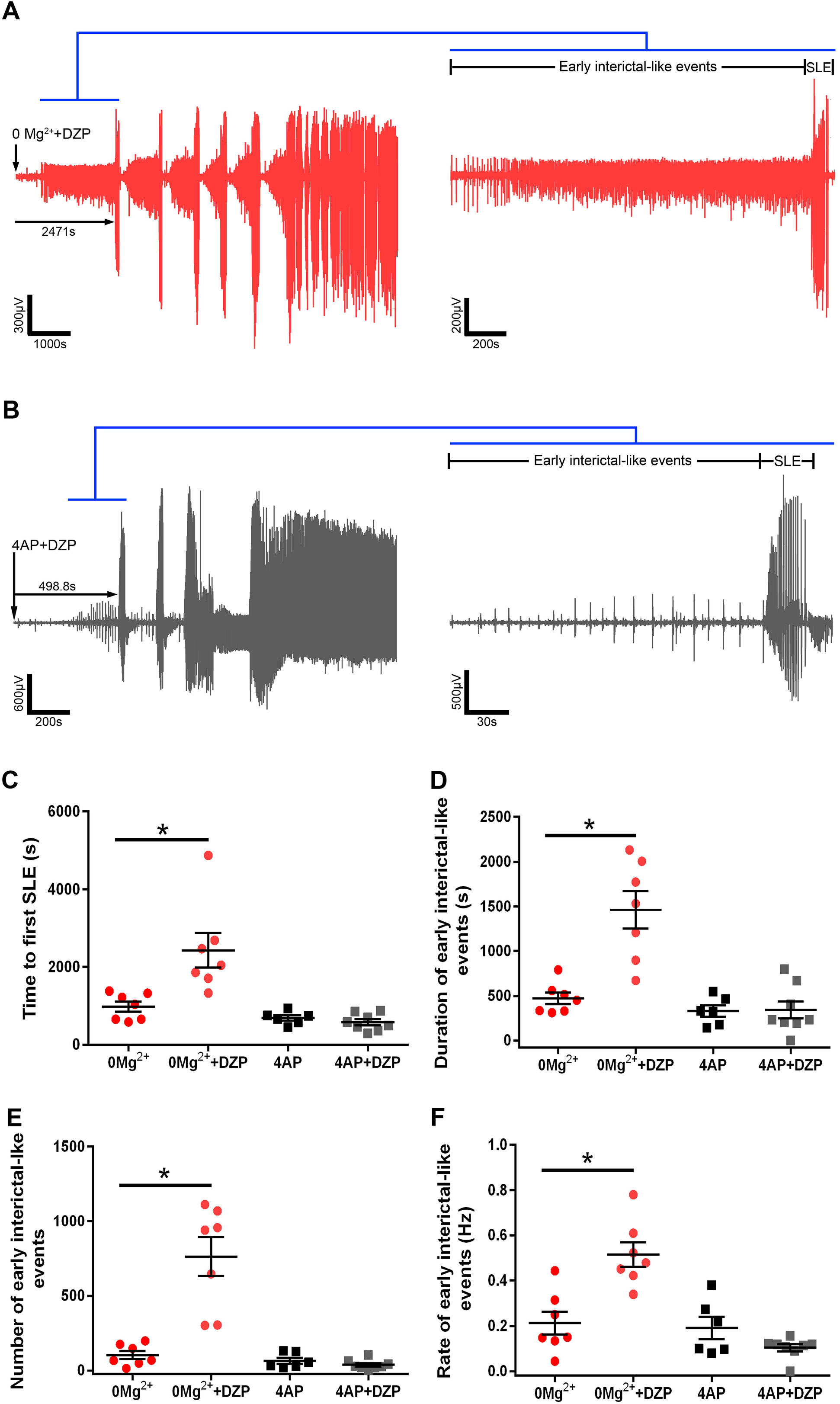
The 0 Mg^2+^ model and the 4AP model differ in their sensitivity to diazepam. A) Representative LFP trace demonstrating the effect of diazepam on 0 Mg^2+^ induced epileptiform activity. B) Equivalent recording in 4AP. C) Diazepam significantly delays seizure-like events in the 0 Mg^2+^ model while have no effect on 4AP induced seizures (ANOVA with post-hoc test, *p = 0.001). D) Diazepam significantly increases the duration of the early interictal period in the 0 Mg^2+^ model while having no effect in 4AP (ANOVA with post-hoc test, *p = 0.001). E) Diazepam significantly increases the number of early interictal-like events in the 0 Mg^2+^ model while this remains unchanged in the 4AP (ANOVA with post-hoc test, *p = 0.001). F) Diazepam significantly increases the rate of early interictal-like events in the 0 Mg^2+^ model while having no effect on the 4AP induced early interictal-like events (ANOVA with post-hoc test, *p = 0.001).

## Discussion

In this study, we made side-by-side comparisons of *in vitro* epileptiform activity induced either by bathing in 0 Mg^2+^ ACSF, or in 100µM 4AP. Brain slices were prepared in the same way (often from the same animals), from young, adult mice. We showed previously that the two models affect hippocampal and neocortical areas differently^32^ (note that the entorhinal cortex appears to align with neocortical activity, rather than the hippocampal circuitry, in this regard), with the hippocampal territories being activated early in 4AP, but very late in 0 Mg^2+^. Here we focused on neocortical activity patterns, and showed that 4AP has an especially strong effect on PV-expressing interneurons (although it also alters, to a lesser extent, activity in both SST-expressing interneurons and pyramidal cells), triggering spontaneous bursting in these cells (similar results have also been reported by Williams and Hablitz^33^). These bursts are readily apparent as isolated short-lasting (<1s) discharges in the LFP. Similar duration discharges are also seen in 0 Mg^2+^ ACSF, and both are commonly referred to by the same term, “interictal events” and we have shown previously that these interictal events involve PV cell firing in both models^34^. A more detailed examination using patch-clamp recordings of pyramidal cells, however, reveal a notable difference between these discharges: the 4AP events are almost purely inhibitory, whereas the 0 Mg^2+^ events also have a large glutamatergic component. This difference is also apparent in the preictal bursts.

It has been reported, previously, that the 4AP *in vitro* model induces low voltage fast onset seizure types^17^. We therefore investigated whether the two models showed different onset patterns, but found instead that both models displayed the low voltage fast onset pattern about 50% of the time, and were just as likely to display a hypersynchronous onset pattern. The variability may be in the proximity to the “ictal focus” within the slice, such that some recordings are of the ictogenic process, whereas others are of the secondary generalization, but the key point is that both models show this heterogeneity. 0 Mg^2+^ seizure-like events showed significantly higher power at frequencies above 60Hz, and far more protracted post-ictal suppression, relative to 4AP events. Indeed, in many slices bathed in 4AP, the interictal discharges resume almost immediately.

We also found a marked difference in the drug sensitivity of the two models: benzodiazepines (Figure 9) have little effect on the evolving activity in 4AP, but markedly reduce the early activity and the time to first seizure-like events in 0 Mg^2+^. Consistent with this, other groups have noted that persistent epileptiform bursting leads to a reduced sensitivity of neurons to diazepam^35–38^. Notably, previous work showed that gap-junction blockers also affect the two models differently, giving mixed results appearing to enhance^39^ activity in 0 Mg^2+^, but suppressing 4AP induced activity^40,41^.

These varied differences between the experimental models provide a powerful research tool for investigating epileptic phenotypes or drug actions in rodent tissue. In transgenic mice carrying mutations associated with epilepsy, it can often be hard to fathom exactly where in the network the problem arises. These acute models though provide a means of “stress-testing” the network^42^, which may help highlight, for instance, whether the problem is primarily neocortical or hippocampal in origin, or to what extent it involves interneuronal dysfunction, and in what way.

A key lesson from these studies is that superficial similarities between epileptiform activity patterns, especially when viewed only using analyses of field potentials, can conceal important differences in the underlying cellular and network activity. This is of obvious clinical importance, since field potentials, most of which are not even ‘local’, but rather surface or scalp EEG, are the only clinical electrophysiological recordings available to us. That is not to say such recordings are worthless, because they clearly do contain huge amounts of information, but we still have much to learn about how to access this for clinical use.

A major motivation behind our study was the important work elucidating ictogenesis in the 4AP model, aided by the development of optogenetic strategies for uniquely activating, or suppressing activity in, specific subpopulations of neurons. Both techniques are compatible with *in vivo* studies, resulting in a large upsurge in the use of 4AP to induce epileptiform activity acutely. It is important therefore to understand how this model gives rise to seizure-like activity, and further, what commonalities, or differences, exist between this model and others. A growing consensus points to a critical role for short-term ionic plasticity affecting particularly fast GABAergic synaptic function (reviewed in ^27^, but see also ^22,26,43,44^). This is exacerbated by other causes of raised intracellular chloride, more chronically^45,46^. On this background of raised [Cl^−^]_i_, bursts of interneuronal activity will excite the most extreme loaded pyramidal cells^33^, and will further load other cells with chloride acutely. Moreover, this raised intracellular Cl^−^ then causes a secondary surge in extracellular [K^+^], due to coupled extrusion of Cl^−^ and K^+^ mediated by KCC2 ^22^.

Specific optogenetic activation of interneurons has been shown to trigger SLEs, albeit not invariably, in 4AP^15,17,18,47^ and in the LRD phase of the 0 Mg^2+^ model^15^. Critically, though, this does not occur early in the 0 Mg^2+^ model^48^, when SLEs still occur, when GABAergic function remains inhibitory and exercises a marked restraint on propagating epileptiform discharges^23^. This restraint is powerful, but also short-lived^23,26,49^, since its enactment provides one of the most rapid means of driving Cl^−^ into neurons, on account of the massive, and concurrent, GABAergic and glutamatergic synaptic drive^26,28,29,49,50^. Thus these early SLEs in the 0 Mg^2+^ also involve a progressively compromised GABAergic function arising from short-term ionic plasticity. This then is the nub of the difference between the two models, that in the absence of appreciable glutamatergic drive, spontaneous or induced interneuronal activity can trigger SLEs through an extreme form of GABAergic dysfunction, involving raised intracellular Cl^−^ and extracellular K^+^. Notably, a recent study of an acute, in vivo model, triggered by direct injection of pilocarpine into neocortex, showed the rapid switch, occurring over a few seconds, of the inhibitory effect of PV-interneuron activation to an excitatory one^51^, consistent with the previous demonstration in organotypic cultures^26^. They further showed that this GABAergic switch could be prevented by over-expressing KCC2 artificially (introduced by viral vectors), providing strong support for the hypothesized role of short-term ionic plasticity in ictogenesis^51^.

Critically, though, in the presence of large glutamatergic drives, ictal recruitment will occur at a far earlier stage in this spectrum of GABAergic dysfunction. It remains an open question for clinical practice which of these two situations is dominant, but one can conceive examples of both. For instance, sensory-triggered seizures, and secondary generalization into cortical areas that had been functioning normally, are both likely to represent cases where the glutamatergic drive is predominant. Optogenetic stimulation of interneurons is an important experimental model, but the key clinical question is whether there are pathological conditions in which one sees spontaneous bursting of interneurons, in the absence of any significant glutamatergic drive. Of course, there could be special clinical cases of this, for instance, such as with genetic mutations of certain classes of K^+^ channels that are expressed predominantly in interneurons^48,52–54^. Any EEG marker of such activity would represent an important means of identifying patients at high risk of having a seizure. Furthermore, a clinical understanding of the source of the epileptiform discharges (presence or absence of appreciable glutamatergic drive) is likely to have relevance for determining the most appropriate pharmacological interventions for individual patients.

## Abbreviations

PV: Parvalbumin
SST: Somatostatin
ACSF: artificial cerebro-spinal fluid
SLE: seizure-like events
LRD: late recurrent discharges

## Author contributions

Experiments and data analysis were performed by NKC, RG, TJ, RJB and RRP. Experiments were conceived by NKC, AJT, JVR, and RRP. Manuscript was written by RRP and AJT, and edited by the other authors. All authors have approved the final manuscript and agree to be accountable for all aspects of the work. All authors qualify for authorship, and all who qualify for authorship are listed.

## Acknowledgements

We would like to thank Meribeth Parrish and Claudia Racca for comments on the manuscript.

